# Telacebec for ultra-short treatment of Buruli ulcer in a mouse model

**DOI:** 10.1101/2020.02.10.943175

**Authors:** Deepak V. Almeida, Paul J. Converse, Till F. Omansen, Sandeep Tyagi, Rokeya Tasneen, Jeongjun Kim, Eric L. Nuermberger

**Affiliations:** Center for Tuberculosis Research, Department of Medicine, Johns Hopkins University, Baltimore, Maryland, USA; Infectious Diseases Unit, Department of Internal Medicine, University of Groningen, Groningen, The Netherlands; Qurient Co. Ltd., Korea

## Abstract

Telacebec (Q203) is a new anti-tubercular drug with extremely potent activity against *Mycobacterium ulcerans*. Here, we explored the treatment-shortening potential of Q203 alone or in combination with rifampin (RIF) in a mouse footpad infection model. The first study compared Q203 at 5 and 10 mg/kg doses alone and with rifampin. Q203 alone rendered most mouse footpads culture-negative in 2 weeks. Combining Q203 with rifampin resulted in relapse-free cure 24 weeks after completing 2 weeks of treatment, compared to a 25% relapse rate in mice receiving RIF+clarithromycin, the current standard of care, for 4 weeks.

The second study explored the dose-ranging activity of Q203 alone and with RIF, including the extended activity of Q203 after treatment discontinuation. The bactericidal activity of Q203 persisted for ≥ 4 weeks beyond the last dose. All mice receiving just 1 week of Q203 at 2-10 mg/kg were culture-negative 4 weeks after stopping treatment. Mice receiving 2 weeks of Q203 at 0.5, 2 and 10 mg/kg were culture-negative 4 weeks after treatment. RIF did not increase the efficacy of Q203. A pharmacokinetics sub-study revealed that Q203 doses of 2-10 mg/kg in mice produce plasma concentrations similar to those produced by 100-300 mg doses in humans, with no adverse effect of RIF on Q203 concentrations.

These results indicate the extraordinary potential of Q203 to reduce the duration of treatment necessary for cure to ≤ 1 week (or 5 doses of 2-10 mg/kg) in our mouse footpad infection model and warrant further evaluation of Q203 in clinical trials.

## Introduction

The World Health Organization’s recommended treatment for Buruli ulcer (BU), also known as *Mycobacterium ulcerans* disease, recently evolved from an 8-week regimen of rifampin (RIF, R) at 10 mg/kg of body weight plus streptomycin (STR) to an 8-week regimen of RIF plus clarithromycin (CLR, C) to eliminate the need for the injectable agent STR and to avoid its related ototoxicity (1, 2). However, CLR has more limited activity than STR against *M. ulcerans* in mouse models of the disease and RIF induces the metabolism of CLR, which likely limits the contribution of CLR to the regimen (3-5). Nonetheless, clinical studies have shown good efficacy of the RIF+CLR regimen (6).

Despite the success of the RIF+CLR regimen, shortening the duration of BU treatment remains an important research objective. We previously investigated replacement of STR and/or CLR with other drugs such as clofazimine and oxazolidinones in our mouse footpad infection model, as well as the impact of increasing rifamycin exposures using high-dose RIF or rifapentine (RPT), with the aim of reducing the treatment duration necessary for cure (7-12). Although we identified novel combinations with efficacy superior to RIF+STR and/or RIF+CLR, none of these 2-drug combinations showed a potential to reduce the duration of treatment to less than 4 weeks in mice.

Telacebec (Q203, Q) is a new drug developed to treat tuberculosis by targeting the respiratory cytochrome bc_1_:aa_3_ complex (13). In *in vitro* and mouse models of tuberculosis, Q203 often exhibits bacteriostatic, rather than bactericidal, activity due to the presence of an alternative terminal oxidase, the cytochrome *bd* oxidase, that maintains electron transport chain (ETC) function and preserves viability (14). However, unlike *Mycobacterium tuberculosis*, classical strains of *M. ulcerans* have a naturally occurring mutation in the *cydA* gene that renders the cytochrome *bd* oxidase non-functional (15). Therefore, most *M. ulcerans* strains causing BU are exquisitely susceptible to Q203 with very low MICs of 0.000075-0.00015 µg/ml (16, 17). *In vivo* studies also show Q203 to be a very attractive candidate for treatment of BU. Scherr et al. (17) showed that Q203 alone at a daily dose of just 0.5 mg/kg was as effective as RIF+STR and rendered 9/10 mice culture-negative with 8 weeks of treatment. Seeking a novel treatment-shortening regimen, we tested Q203 at 10 mg/kg/day in 3- and 4-drug combinations with high-dose RPT and other drugs acting on the ETC and oxidative phosphorylation (clofazimine (CFZ), and bedaquiline (BDQ)) and found that mouse footpads were sterilized after just 2 weeks of treatment (16).

In the present work, we explored the treatment-shortening potential of simpler, more readily implementable regimens based on Q203 alone or in combination with RIF. Two sequential experiments in the mouse footpad infection model assessed the dose-ranging efficacy of Q203 with or without normal and high-dose rifampin in tandem with pharmacokinetics (PK) analysis to better understand the human-equivalent doses. The results demonstrate that Q203 exposures recently demonstrated in phase 1 trials (18), are capable of sterilizing mouse footpads after as little as 1 week of treatment (5 doses), making Q203 an extraordinary candidate for clinical trials to shorten BU treatment.

## Results

### Study 1: To determine the sterilizing efficacy of combining Q203 with standard and high doses of rifampin

To determine if replacing CLR with Q203 in the RIF+CLR regimen has the potential to shorten the treatment of BU, we assessed the sterilizing efficacy of Q203 at 5 or 10 mg/kg when combined with RIF at 10 and 20 mg/kg doses.

#### Footpad swelling and CFU counts

Mean (± SD) footpad CFU counts on the day after infection were 2.71 ± 0.93 log_10_ CFU/footpad. Six weeks later, at the start of treatment (D0), the median swelling grade was ≥ 2.5 on a scale of 0-4 (10, 19) (Fig. 1) and the mean CFU count reached 5.42 ± 0.56 log_10_ CFU/footpad (Fig. 2). After 1 week of treatment, all treatment groups receiving Q203 had markedly reduced footpad swelling compared to R_10_C_100_ controls (Fig. 1) (hence forth drugs are denoted in single letters followed by the dose in mg/kg shown in subscripts). The swelling grade in the R_10_C_100_ group remained unchanged, while mice treated with Q203-containing regimens, with exception of Q_5_ alone, all had medians of ≤1 (Fig. 1). Similarly, the CFU counts at week 1 in the R_10_C_100_ group were significantly higher than those in all RQ groups except the Q_10_ alone group (Fig. 2A). After 2 weeks of treatment, footpads in all Q203-treated groups were almost normal compared to the R_10_C_100_ group which still had swelling with a median grade of 2. The corresponding footpad cultures were negative in all R_20_Q_10_-treated mice and negative in nearly all other Q203-treated groups compared to 2.63 ± 0.37 log_10_ CFU in the R_10_C_100_ group (Fig. 2B). The limit of detection was 3 CFU per footpad. After 4 weeks of treatment, all mice had normal footpads and no CFU were detected in any of the treatment groups tested. The limit of detection was 1 CFU per footpad. Mean CFU counts are provided in supplementary Table S1.

**Fig 1.**
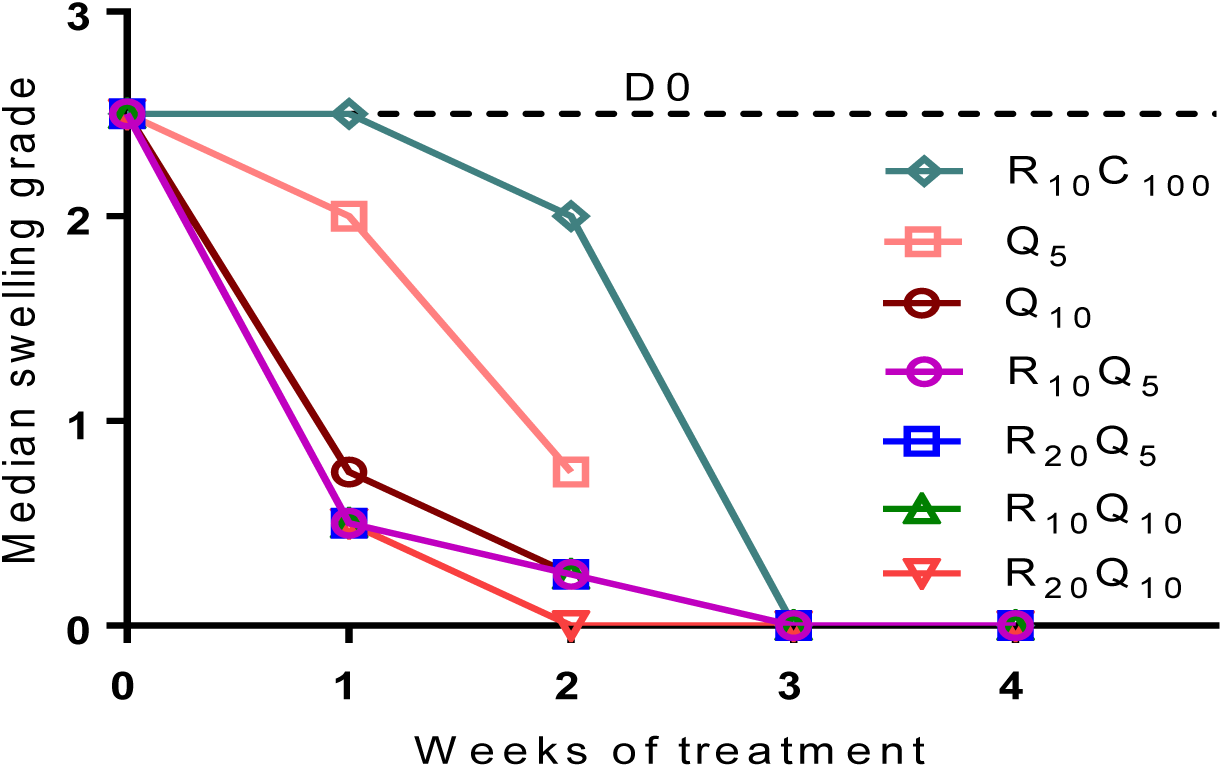
Footpad swelling grade of infected mouse footpads in response to treatment in Study 1: Treatment was initiated 6 weeks after infection, when swelling approached swelling grade 2.5. Swelling grade 0 corresponds to no clinically visible pathology, grade 1 infers redness of the footpad, grade 2 edematous swelling of the footpad, and grade 3 ascending swelling of the leg and impeding necrosis. Data points represent medians per treatment group. Data were normalized to day 0 (beginning of treatment) by subtracting from the median swelling grade of all mice at D0 and assuming the total median as group mean for that time point. All Q-containing regimens rapidly reduced swelling grade compared with R_10_C_100_ controls. By the end of 1 week of treatment, all Q-containing regimens, except the lowest dose of Q, 5 mg/kg, had reduced the swelling to below grade 1, while no change was seen in the RC treatment controls. By the end of 2 weeks in all mice treated with Q-containing regimens had only residual swelling left, median swelling grade was 0.25. Numbers in subscripts after drugs indicate doses in mg/kg. D, day; R, rifampin; C, clarithromycin; Q, Q203/Telacebec.

**Fig 2.**
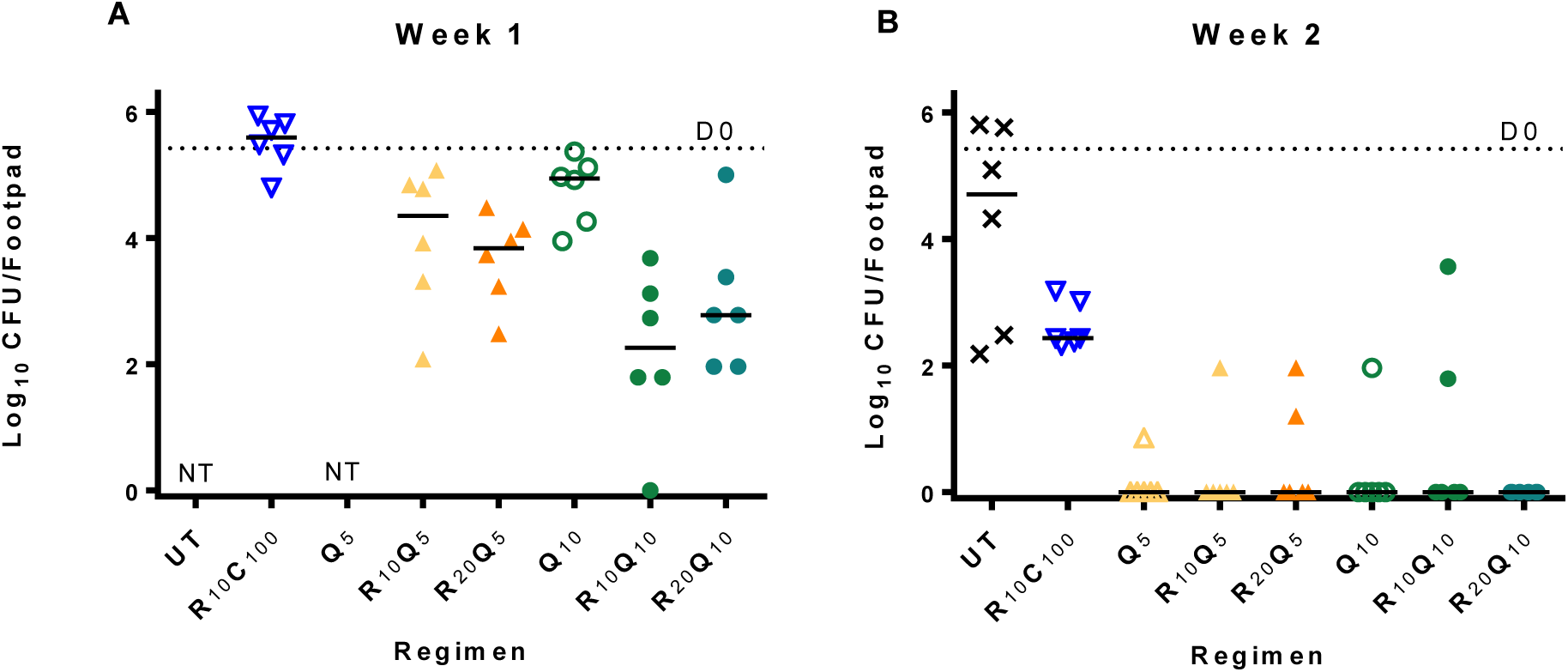
Microbiological outcome in Study 1: Mice were infected with 2.71 ± 0.93 log_10_ CFU/footpad of *M. ulcerans* into both hind footpads. After 6 weeks of incubation, treatment was initiated (D0). At this time point, the CFU mean (±SD) equaled 5.42 (±0.56). Groups of mice were sacrificed at week 1, week 2 and week 4 and footpads (n= 6) were dissected, minced, and plated on 7H11 selective agar for colony counting and CFU analysis. For statistical analysis all test regimens were compared to R_10_C_100_ controls. (A) After 1 week of treatment all Q-containing regimens, except Q_10_ given alone, were significantly better than controls, with R_10_Q_10_ (p < 0.0001) and R_20_Q_10_ (p < 0.001) showing the best activity. (B) At week 2, most footpads in mice treated with Q-containing regimens were culture negative and significantly better than R_10_C_100_ (p ≤ 0.0008). At week 4, none of the mice in the combination treatment groups, including R_10_C_100_ controls, were culture-positive (data not shown). Monotherapy regimens were not tested at this timepoint. Numbers in subscript after drugs indicate doses in mg/kg. D, day; UT, untreated; R, rifampin; C, clarithromycin; Q, Q203/Telacebec; NT, Not tested. Dashed line indicates the pre-treatment CFU at D0. Horizontal lines indicate median values.

#### Relapse

Relapse assessments were made 6 months after treatment completion in mice treated for 2 or 4 weeks. All mice treated with RIF+Q203 regimens showed no rebound in footpad swelling during the 6-month follow-up period and the CFU counts were all zero (limit of detection: 1 CFU). In the R_10_C_100_ group relapse was assessed only in mice treated for 4 weeks. Three mice experienced a rebound in footpad swelling during the 6-month follow-up period. Two of these mice required euthanasia before the planned relapse endpoint because one or both footpads had deteriorated beyond a lesion index of 3. Overall, the relapse rate in the R_10_C_100_ group was 25%, with 4/16 footpads positive for CFU (p=0.10 vs. other groups with 0/16 relapses).

### Study 2: To determine the dose-ranging activity of Q203 alone and in combination with standard and high-dose rifampin, including the extended activity after treatment discontinuation

After showing that Q203 alone at 5-10 mg/kg/day renders mouse footpads culture-negative and combinations of RIF+Q203 sterilize footpads with just 2 weeks of treatment, we evaluated a lower dose range and shorter durations of Q203, alone and in combination with RIF. We also assessed the plasma PK of Q203 after single doses of 0.5, 2 and 10 mg/kg doses, and determined plasma concentrations of Q203 at 3-4 days and 2 and 4 weeks after stopping treatment.

#### Pharmacokinetics

The single dose PK results for Q203 are shown in Fig. 3. Q203 had a t_max_ of 2 hrs. C_max_ values after 0.5, 2 and 10 mg/kg doses were 0.05, 0.17 and 0.92 µg/ml respectively, indicating that even at the lowest dose, the C_max_ was well above the MIC of 0.000075-0.00015 µg/ml. Plasma AUC values indicated dose-proportional exposures up to 2 mg/kg. After 1 week of treatment, in the lowest dose group tested, Q203 at 0.5 mg/kg, the mean plasma concentration of Q203 at 72 hrs post-dose was 0.073 ± 0.024 µg/ml and in the groups treated for 2 weeks, at 96 hrs post-dose, it was 0.052 ± 0.013 µg/ml (Supplementary Table S2). In both these groups, the concentrations gradually declined during the 4-week follow-up period, but remained higher than the MIC. With higher doses of Q203, more accumulation was seen, and it increased with the duration of treatment, with plasma concentrations in mice treated for 2 weeks almost twice as high as in those treated for one week. In mice treated with RIF+Q203, Q203 concentrations were similar to those in mice treated with Q203 alone indicating no large effect of RIF on Q203 concentrations.

**Fig 3.**
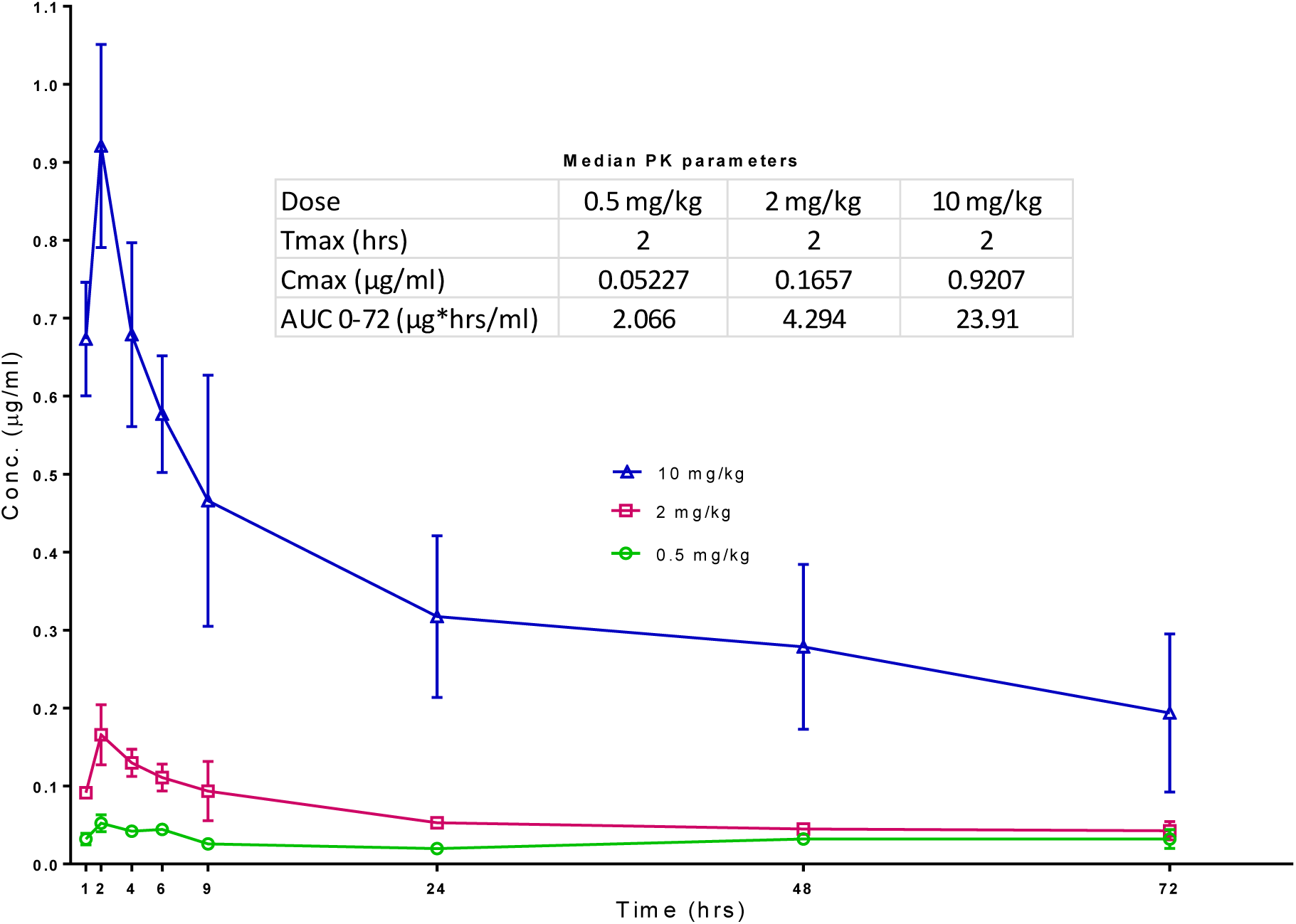
Single dose pk for Q203: Mice were dosed with either 0.5 mg/kg (green circle), 2 mg/kg (red squares) or 10 mg/kg (blue triangles) dose of Q203 and the blood collected for serum concentrations at the indicated timepoints. Median PK parameters shown in the inset indicate dose-proportional exposures.

#### Footpad swelling

At the start of treatment, mice had a median swelling grade of 2 (Fig. 4). In untreated mice, the swelling increased to grade 2.5 and grade 3 at end of Weeks 1 and 2, respectively. All untreated mice required euthanasia at this point. In mice treated with R_10_C_100_, there was a marginal decrease to 1.5 at Week 1 (Figs. 4A and 4B) and a further decrease to 0.5 at Week 2 (Fig. 4 C and D). After stopping treatment at 2 weeks, the swelling continued to decrease gradually during the 4 weeks of follow-up and reverted to baseline (Figs. 4C and D). In mice treated with RIF 10 mg/kg alone for 1 week, there was slight decline in swelling after peaking at week 1 (Fig. 4A). At the end of 4 weeks, when all mice were sacrificed for CFU, the median swelling grade was 1.5, with one mouse showing grade 3 swelling. In mice treated with RIF 10 mg/kg for 2 weeks (Fig. 4C), the response to treatment was better than in mice treated for only 1 week. After 2 weeks of treatment, the swelling had reduced to median swelling grade of 1.5 and gradually decreased to 0.5 after 4 more weeks of follow-up without treatment. In mice treated with RIF 20 mg/kg, the response to treatment was similar to that of 10 mg/kg group. As in the previous experiment, footpad swelling decreased rapidly in Q203-treated groups. All doses of Q203 whether given alone or in combination with RIF at 10 or 20 mg/kg rapidly reduced footpad swelling. After 1 week of treatment, the swelling reverted to baseline in most mice, while some mice showed residual swelling (swelling grade < 1) (Fig. 4A). Irrespective of whether the treatment was stopped after one week or continued for an additional week, the footpads continued to improve during the follow-up period, with almost all footpads returning to baseline by Week 2 and remaining free of swelling.

**Fig 4.**
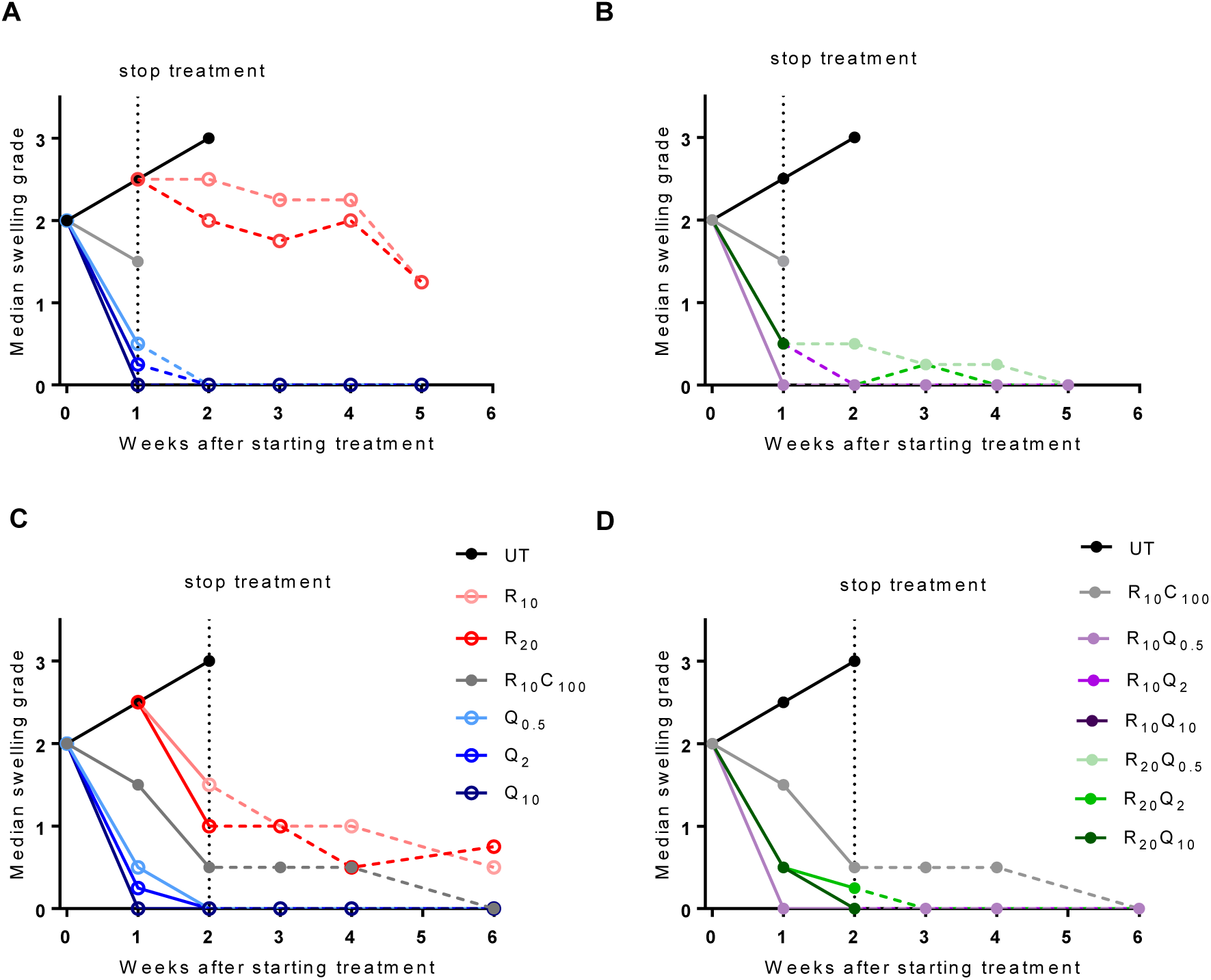
Footpad swelling grade of infected mouse footpads in response to treatment in Study 2: Treatment was initiated 6 weeks after infection, when median swelling grade approached 2. Data points represent medians per treatment group. Swelling results in mice treated for 1 week are shown in panels A and B and those for mice treated for 2 weeks are shown in panels C and D. Monotherapy groups are shown in panels A and C while combination treatment groups are shown in panels B and D. R_10_C_100_ is the standard treatment control. Solid lines represent change in footpad swelling during treatment, while that after stopping treatment is shown by dashed line. All Q-containing regimens reduced footpad swelling after just 1 week of treatment, and continued to show response after stopping treatment. Most footpads were at baseline levels after 2-3 weeks. R alone produced a slight decline in swelling after peaking at 1 week. The footpads never reached a median grade of 1 after 4 weeks follow-up. As with 1 week of treatment, 2 weeks of Q-containing regimens rapidly rendered footpads swelling-free. In comparison, the RC-treated controls showed gradual decreases in footpad swelling. Numbers in subscripts after drugs indicate doses in mg/kg. D, day; UT, untreated; R, rifampin; C, clarithromycin; Q, Q203/Telacebec,

#### Footpad CFU counts

At the start of treatment (D0), mean footpad CFU counts were 5.40 ± 0.39 log_10_. They increased to 5.71 ± 0.23 at Week 1 (Figs. 5A and 5B) and 5.98 ± 0.23 at Week 2 in untreated mice (Figs. 5C and 5D).

**Fig 5.**
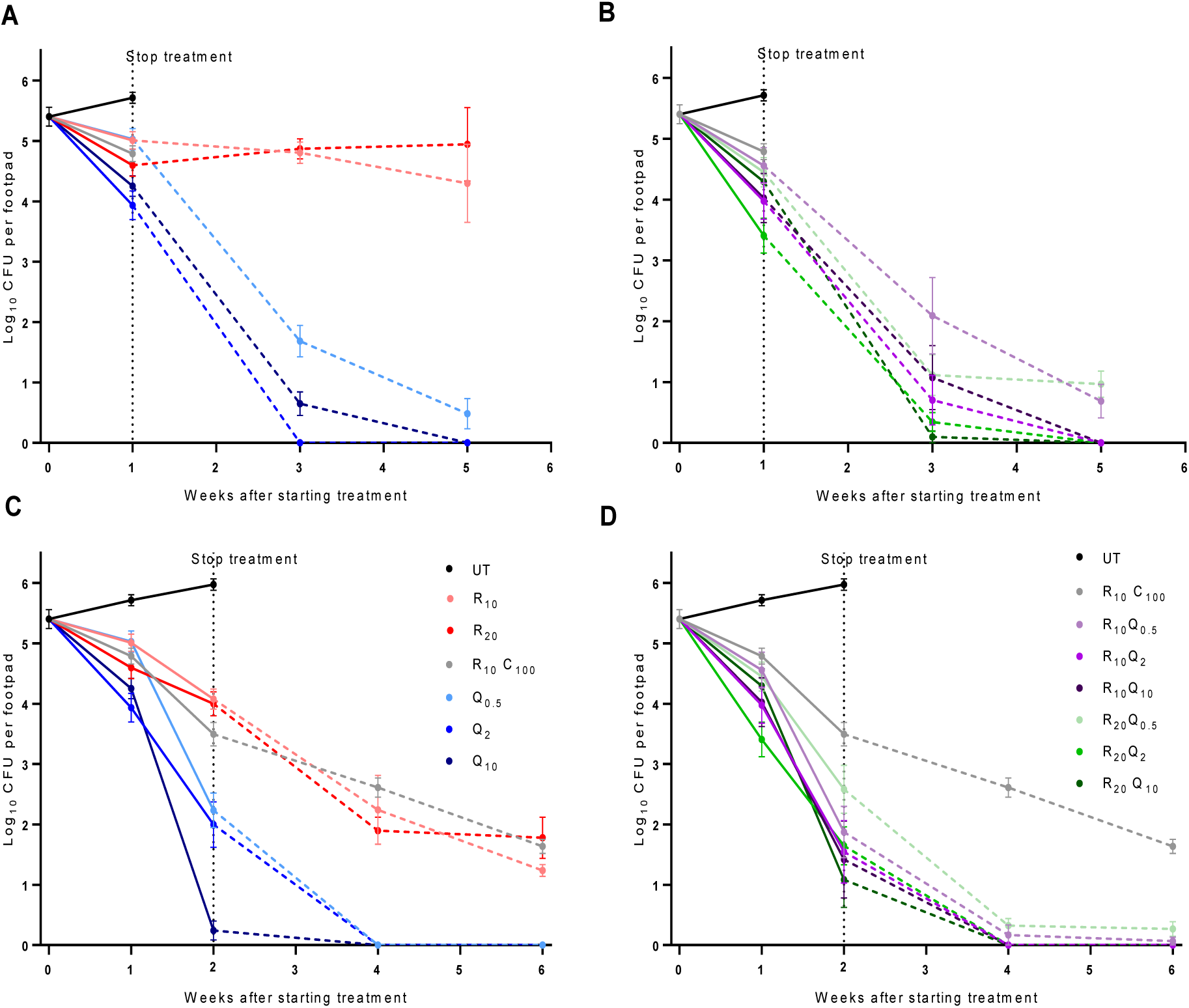
Microbiological outcome in Study 2: Panels A and B show the response to treatment for 1 week, and panels C and D show the response to treatment for 2 weeks. Panels A and C show results for monotherapy, and panels B and D show combination treatment groups. Solid lines indicate fall in mean CFU (±SEM) during treatment and dashed line shows reduction after stopping treatment.. After 1 week of treatment, Q-containing regimens showed a marked dose response, and although CFU counts at Week 1 were not significantly different than RC controls, more dramatic reductions occurred during the 4 week follow-up period after stopping treatment. All Q-containing regimens except Q_0.5_ were significantly better after 1 week of treatment than RC treatment for 2 weeks. After 2 weeks of treatment, all Q-containing regimens were significantly better than RC control after 2 weeks and rendered footpads negative at follow-up 2 weeks after stopping treatment.

In mice treated with R_10_C_100_, CFU counts decreased to 4.79 ± 0.32 log_10_ at Week 1 and 3.50 ± 0.48 log_10_ at Week 2. Continued killing was observed after stopping treatment, corroborating the observed reductions in footpad swelling during the 4-week follow-up period (Figs. 5C and 5D). In mice receiving RIF at 10 or 20 mg/kg, little change in CFU was seen after one week of treatment or during the 4-week follow-up period, again similar to what was seen in footpad swelling (Figure 5A). Increasing the duration of RIF treatment to two weeks resulted in a 1.5 log_10_ CFU reduction in both dosage groups (Fig. 5C) and continued decreases to 1.24 ± 0.23 and 1.78 ± 0.83 4 weeks after completing treatment in the R_10_ and R_20_ groups, respectively.

A modest dose-dependent effect was observed in mice treated with Q203 alone for one week, with CFU counts falling to 5.03 ± 0.43, 3.93 ± 0.58, and 4.25 ± 0.42 log_10_ CFU in those receiving 0.5, 2 and 10 mg/kg doses, respectively. With the exception of R_20_Q_2_ (Fig. 5B) (p= 0.002), the reductions in CFU at Week 1 were not significantly better than the R_10_C_100_ control. However, Q203 treatment for one week resulted in more dramatic reductions in CFU counts after treatment cessation. CFU counts in Q_0.5_-treated mice fell to 1.69 ± 0.64 and 0.48 ± 1.48 log_10_ CFU after 2 and 4 weeks of follow-up. No CFU were detected after 2 or 4 weeks of follow-up in mice treated with Q_2_ or 4 weeks after treatment with Q_10_ (Fig. 5A). Comparisons to the R_10_C_100_ group at W1+2 and W1+4 time points was not possible since we did not include these time points for this group. However, all groups receiving Q203 doses ≥ 2 mg/kg had significantly lower CFU counts at the W1+2 (p ≤ 0.0006) and W1+4 (p < 0.0001) time points than R_10_C_100_ controls had at the W2+2 and W2+4 time points, respectively.

In mice treated for 2 weeks with Q203, a more prominent dose-response relationship was observed. In mice treated with Q203 at 10 mg/kg, the mean log_10_ CFU count was only 0.24 ± 0.38, with 4/6 pads negative. No CFU were detected in footpads in any Q203-treated group at 2 and 4 weeks follow-up (Fig. 5C). At this point all Q203-containing regimens were significantly better when compared with the R_10_C_100_ controls (p ≤ 0.05).

No benefit of adding RIF at either 10 or 20 mg/kg was seen, as CFU counts were very similar to those in groups receiving Q203 alone. In fact, addition of RIF may have been slightly antagonistic at later time points, especially after 2 weeks of treatment and follow-up time points (Figs. 5B and 5D). Mice in the R_10_Q_2_ group received slightly more Q in the first week due to accidental gavage of mice with 10 mg/kg on Day 4. This group was not treated on Day 5 and thus received a 16 mg/kg total dose for the weeks as opposed to the intended 10 mg/kg. By the end of 2 weeks of treatment they had received a 26 mg/kg total dose rather than the intended 20 mg/kg total dose. Mean CFU counts are given in Supplementary Table S3.

## Discussion

The current treatment for BU recommended by WHO (1) is an oral regimen of RIF+CLR given daily for 8 weeks. This regimen offers advantages over the previously recommended RIF+STR combination (2). However, it remains problematic because treatment duration inversely correlates with adherence and patients are often hospitalized until there is clear-cut evidence of efficacy treatment response, including resolution of any paradoxical reaction (20), resulting in missed school or work activities. As an extremely potent inhibitor of *M. ulcerans* respiration, Q203 is an exceptional candidate for treatment-shortening regimens. Recently, we described 3-drug combinations of drugs active on the ETC with and without rifapentine that appeared capable of shortening the treatment of BU (16). Q203-containing regimens proved to be most effective and cured all mice after treatment for just 2 weeks. However, none of the companion drugs in those regimens is currently used in the treatment of BU. Reasoning that RIF is already a core component of BU treatment regimens, we aimed to test Q203 alone and in combination with RIF, comprising regimens easier to implement in the clinical setting. Our results show that regimens of Q203 alone or in combination with RIF are clearly superior to RIF+CLR and may be capable of reducing the duration of BU treatment to 1-2 weeks.

Little information about the PK of Q203 in humans exists in the public domain (18) and published PK data from mice report very different drug exposures for the same or similar doses (13, 17). To better understand the dose-response profile of Q203 in mice and the human equivalent doses of the Q203 doses evaluated in our model, we evaluated Q203 doses ranging from 0.5 to 10 mg/kg and included PK analyses. Remarkably, Q203 alone at 2 mg/kg rendered mouse footpads culture-negative after just 5 daily doses. The median plasma C_max_ and AUC_0-72h_ values after a single oral dose in mice were 0.17 µg/ml and 4.3 µg-h/ml, respectively. These results are in line with mouse PK results from Pethe et al (13) and, more importantly, comparable to the plasma C_max_ and AUC_0-inf_ values of 0.38 µg/ml and 6.3 µg-h/ml, respectively, after a single dose of 100 mg in fed human (18) subjects. Considering that we observed dose-proportional PK in mice, Q203 doses of 2-10 mg/kg in mice likely correspond well to the daily doses of 100-300 mg that were recently reported to be well tolerated and safe in phase 1 trials and in TB patients over 14 days of dosing in a recent phase 2a trial (21), provided that the drug is administered with food. Therefore, we predict that these doses can safely shorten treatment of BU to 5 doses or less.

The extreme treatment-shortening effects of Q203 we observed in mice were the result of persistent killing of *M. ulcerans* that extended well beyond the end of dosing, even at the lowest dose of 0.5 mg/kg. This persistent killing is likely a function of multiple phenomena: exceptionally potent activity (e.g., very low MIC), low clearance (e.g., long plasma half-life), favorable partitioning into tissue (e.g., lung:plasma concentration ratio of 2-3) (13), and a post-antibiotic effect (e.g., continued antimicrobial effect against *M. ulcerans* after plasma concentrations fall below MIC). While the roles of the first 2 phenomena are self-evident from the PK/PD data generated in Study 2, more evidence is needed to confirm the partitioning of Q203 into mouse footpads and presence of a post-antibiotic effect. Treatment with 0.5 mg/kg for 1-2 weeks resulted in persistent killing for at least 4 weeks beyond the end of dosing. Although plasma concentrations were approximately 5-10 times higher than MIC at 2 weeks post-treatment and in the MIC range at 4 weeks post-treatment, Q203 is 99.8% protein bound in mouse plasma. Therefore, it seems likely that free drug concentrations at the site of infection fell below MIC during the 4-week follow-up period, thus suggesting the presence of a post-antibiotic effect. *M. ulcerans* may be especially vulnerable to post-antibiotic effects *in vivo* if drug treatment shuts down production of the immunosuppressive mycolactone toxin, allowing a more effective host immune response to develop and enhance bacterial clearance. Indeed, even RIF+CLR exhibited persistent effects after the end of dosing in Study 2.

Another surprising finding of these experiments was that the addition of RIF did not significantly increase the treatment efficacy of Q203. In fact, other than some additive effects of RIF with Q203 at 0.5 mg/kg at the Week 1 and Week 2 time points, there were hints of modest antagonistic effects of RIF at each Q203 dose level, especially 2 mg/kg and above. Our PK results did not show any significant differences in Q203 plasma concentrations when RIF and Q203 were co-administered when compared to Q203 given alone. These results raise the prospect of using Q203 as monotherapy, a scenario that may be defensible because the spontaneous frequency of Q203 resistance mutations in *M. ulcerans* appears to be very low (17) and *M. ulcerans* is not transmitted from person-to-person, making resistance development both unlikely to occur as well as unlikely to have any impact beyond the affected individual.

In summary, we have demonstrated the extraordinary potential of Q203 to reduce the duration of treatment for BU to 1 week (or 5 doses of 2-5 mg/kg) in our mouse footpad infection model. As these doses appear to be a good representation of doses recently tested successfully in humans, they warrant consideration for further evaluation in clinical trials for BU treatment. Importantly, we did not define the shortest duration of Q203 treatment needed to eradicate *M. ulcerans* from mouse footpads. Studies evaluating even shorter durations of Q203 with and without additional companion drugs are underway.

## Methods and Materials

### Bacterial strain

*M. ulcerans* strain 1059, originally obtained from a patient in Ghana, was used for the study (22).

### Antibiotics

RIF was purchased from Sigma. CLR was purchased from the Johns Hopkins Hospital pharmacy. Q203 was kindly provided by the Global Alliance for TB Drug Development. RIF and CLR were prepared in sterile 0.05% (wt/vol) agarose solution in distilled water. Q203 was formulated in 20% (wt/wt) D-α tocopheryl polyethylene glycol 1000 (Sigma) succinate solution.

### Mouse infection

BALB/c mice (Charles River Laboratories) were inoculated subcutaneously in both hind footpads with 0.03 ml of a culture suspension containing *M. ulcerans* 1059. Treatment began 6-7 weeks (D0) after infection when the mice had footpad swelling of grade ≥2.

### Treatment

Mice were treated 5 days per week in 0.2 ml by gavage. Drug doses were chosen based on mean plasma exposures (i.e., similar area under the concentration-time curve over 24 hours post-dose in blood) compared to human doses (7, 16). All animal procedures were conducted according to relevant national and international guidelines and approved by the Johns Hopkins University Animal Care and Use Committee.

#### Study 1

Mice were randomized to one of the seven treatment groups (Supplemental Table S4). Control regimens included R_10_C_100_, Q_5_ alone or Q_10_ alone, where the subscript represents the dose in mg per kg of body weight. Test regimens consisted of either R_10_Q_5_, R_20_Q_5_, R_10_Q_10_ or R_20_Q_10_, and mice were treated for either 2 or 4 weeks. Mice treated with Q alone were treated for only 2 weeks since they were only included in the experiment to inform the contribution of Q203 and we did not initially intend to explore the use of Q203 alone as monotherapy. CFU counts were performed after 1, 2 and 4 weeks of treatment to determine the response to treatment. To determine the sterilizing activity of each test regimen, mice were held without treatment for six months after completing 2 and 4 weeks of treatment. Relapse assessment for the R_10_C_100_ control group was done only after 4 weeks of treatment.

#### Study 2

Mice were randomized to one of 12 treatment groups, which included Q203 at doses of 0.5, 2 and 10 mg/kg given alone or in combination with RIF at 10 or 20 mg/kg. Control groups were untreated or received R_10_ or R_20_ alone, or R_10_C_100_, which is the current standard of care (Supplemental Table S5). Mice were treated for either 1 or 2 weeks. R_10_Q_2_ group mice were accidentally gavaged with Q at 10 mg/kg dose on Day 4 of treatment. These mice were not gavaged on the following day and therefore received a cumulative dose of 16 mg/kg instead of the intended 10 mg/kg dose for the first week. By the end of 2 weeks of treatment, these mice had received a total dose of 26 mg/kg instead of the intended 20 mg/kg dose. Footpad CFU counts were done at treatment completion and also at 2 and 4 weeks after stopping treatment in each treatment group to determine the continued bactericidal activity of Q203-containing regimens.

### Pharmacokinetics

Intensive PK evaluation was done for groups receiving Q203 alone and in combination with RIF. After a single dose on D0, small-volume blood samples were collected in EDTA-containing tubes at 1, 2, 4, 6, 9, 24, 48 and 72 hrs post-dose from the submandibular vein. To assess the clearance of Q203 after stopping treatment, samples were obtained at 72 hrs after the final dose in mice treated for 1 week and 96 hrs after the final dose in mice treated for 2 weeks. Blood samples were also obtained at the 2- and 4-week follow time points in mice treated with Q203 alone for 1 and 2 weeks, and in mice treated with RIF+Q203 for 2 weeks.Samples from the R_10_Q_2_ group were excluded because mice in this group were accidentally gavaged with 10 mg/kg dose on Day 4 of treatment.

### Evaluation of treatment response

Treatment outcomes were evaluated based on (i) decrease in footpad swelling, denoted as swelling grade, and (ii) decrease in CFU counts. The swelling grade was was scored as described previously (7). Briefly, the presence and the degree of inflammatory swelling of the infected footpad were assessed weekly and scored from 0 (no swelling) to 4 (inflammatory swelling extending to the entire limb) for all surviving mice. For CFU counts, six footpads (from three mice) were evaluated on the day after infection (D-42), and at the start of treatment (D0) to determine the infectious dose and the pretreatment CFU counts, respectively.

The response to treatment was determined by plating 6 footpads (from 3 mice) from each treatment group at predetermined time points. Footpad tissue was harvested after thorough disinfection with 70% alcohol swabs and then homogenized by fine mincing before suspending in sterile phosphate buffered saline (PBS). Ten-fold serial dilutions and undiluted fractions of homogenate were plated in 0.5 ml aliquots on selective 7H11 agar and incubated at 32°C for up to 12 weeks before CFU were enumerated. In the second study, homogenates were plated on 7H11 agar supplemented with 10% OADC and 5% bovine plasma albumin to reduce any potential effects of Q203 carryover due to its long half-life (23).

To determine the sterilization activity of each test regimen in Study 1, mice were held for relapse assessment for 6 months after completing 2 and 4 weeks of treatment. Results were compared to those from mice treated with R_**10**_C_**100**_ for 4 weeks. Footpads were inspected every 2 weeks for any signs of re-swelling after stopping treatment. When re-swelling was observed, mice were sacrificed when the swelling reached a lesion index ≥ 3 and the footpads were harvested and plated for CFU counts. At the end of the 6-month follow-up period, all remaining mice were sacrificed and their footpads (16 footpads in each group) were harvested and plated for CFU. Study 2, instead of relapse assessment at 6 months, we held mice without treatment for an additional 2 or 4 weeks after treatment completion before harvesting and plating for determination of CFU counts.

### Statistical analysis

GraphPad Prism 6 was used to compare mean CFU counts in Q203-containing groups to the R_10_C_100_ control group using two-way analysis of variance with Bonferroni’s post-test to adjust for multiple comparisons. Proportions were compared using Fisher’s Exact test.

## Acknowledgments

This study was supported by the National Institutes of Health (R01-AI113266). We gratefully acknowledge TB Alliance for providing Q203 and Qurient for quantifying Q203 in mouse plasma. We thank Dr Kingsley Asiedu for fruitful discussions and encouragement at the beginning of this study.

## Supplementary Data

**Fig S1.**
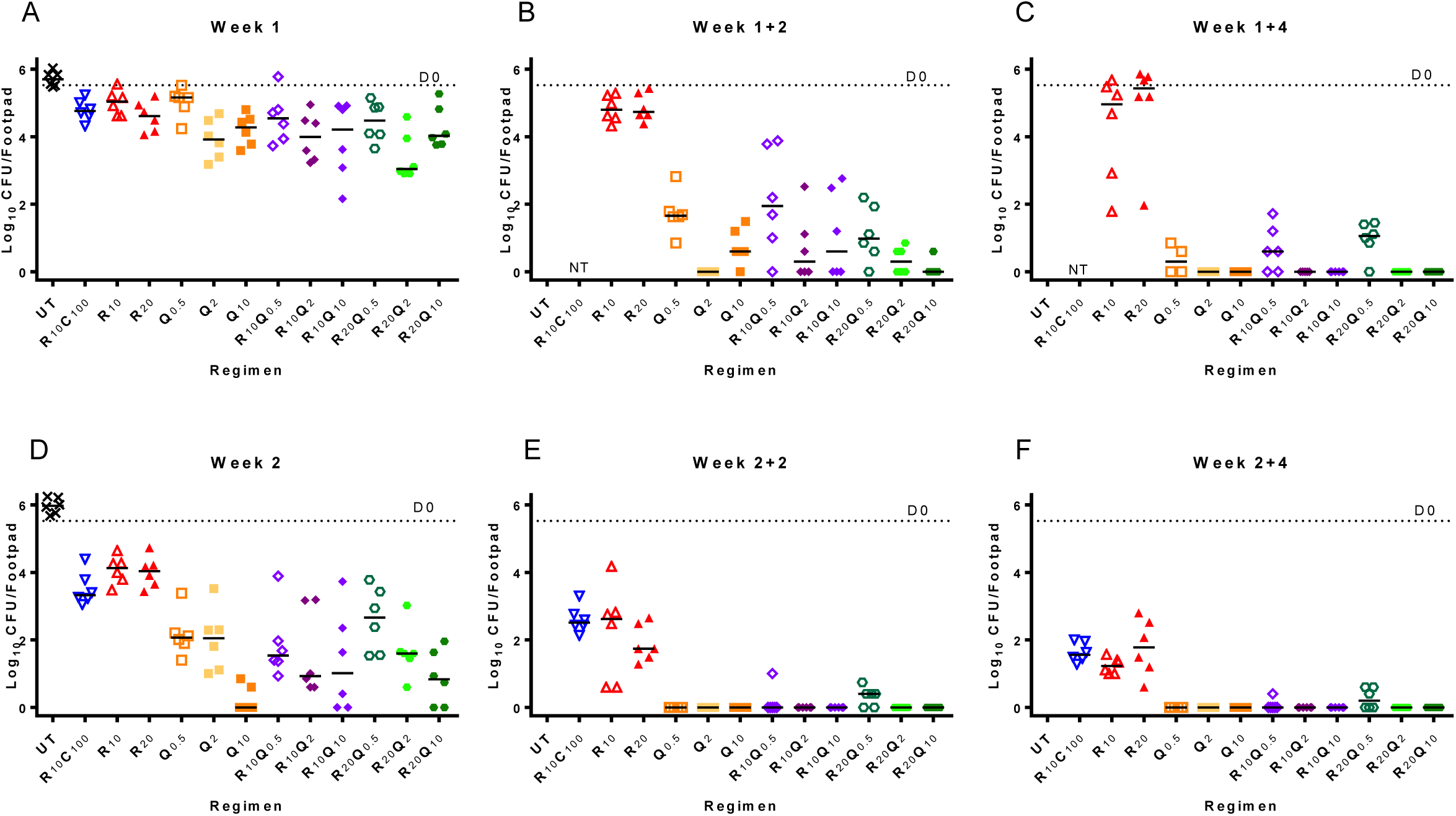
Individual mouse CFU data from Study 2: At the start of treatment (D0) the log_10_CFU/footpad was 5.4 ± 0.39. R_10_C_100_ was the standard treatment control. Top panels show CFU counts in mice treated for 1 week, while bottom panels show CFU counts in mice treated for 2 weeks. (A) After 1 week of treatment, there was not much reduction in CFU in R_10_, R_20_ and R_10_C_100_ controls. Q-treated groups had marginally lower CFUs that were not significantly different from controls, except for R_20_Q_2_ (P = 0.002). (B) At follow-up 2 weeks after stopping treatment (Week 1+2), CFU counts continued to decrease in Q-containing arms. (C) At follow-up 4 weeks after stopping treatment, all mice treated with Q-containing regimens, except those treated with Q at 0.5 mg/kg, were culture-negative. (D) After 2 weeks of treatment, all Q-containing regimens were significantly better than R_10_C_100_ control. (E and F) CFU counts continued to decrease at follow-up 2 and 4 weeks after stopping treatment, with all Q-containing regimens rendering footpads culture-negative.

**Table S1:** Change in footpad CFU in Study 1: *To determine the sterilizing efficacy of combining Q203 with standard and high doses of rifampin*.

| Regimen | Mean (±SD) CFU count | | | | | Proportion of footpads with positive culture | |
| --- | --- | --- | --- | --- | --- | --- | --- |
|  | D-40 | D0 | Week 1 | Week 2 | Week 4 | Week 2 (+24) | Week 4 (+24) |
| <u>Controls</u> |  |  |  |  |  |  |  |
| Untreated | 2.71 ± 0.93 | 5.42 ± 0.56 | | 5.24 ± 0.69 | | | |
| R<sub>10</sub>C<sub>100</sub> | | | 5.51 ± 0.41 | 2.63 ± 0.37 | 0 | NT | 4/16 |
| Q<sub>5</sub> | | | NT | 0.14 ± 0.35<sup>1</sup> | NT | | |
| Q<sub>10</sub> | | | 4.77 ± 0.54 | 0.33 ± 0.80<sup>1</sup> | NT | | |
| <u>Tests</u> |  |  |  |  |  |  |  |
| R<sub>10</sub>Q<sub>5</sub> | | | 4.00 ± 1.15 | 0.33 ± 0.80<sup>1</sup> | 0 | 0/16 | 0/16 |
| R<sub>20</sub>Q<sub>5</sub> | | | 3.67 ± 0.72 | 0.53 ± 0.85<sup>2</sup> | 0 | 0/16 | 0/16 |
| R<sub>10</sub>Q<sub>10</sub> | | | 2.18 ± 1.30 | 0.89 ± 1.49<sup>2</sup> | 0 | 0/16 | 0/16 |
| R<sub>20</sub>Q<sub>10</sub> | | | 2.98 ± 1.13 | 0 | 0 | 0/16 | 0/16 |

<sup>1</sup> 5/6 pads negative
<sup>2</sup> 4/6 pads negative

Day after infection is D-40, D0 is the start of treatment. Relapse was assessed 24 weeks after completing 2 weeks (2+24) and 4 weeks (4+24) of treatment. Drug abbreviations: R, Rifampin; C, Clarithromycin; Q, Q203/Telacebec. Drug dose is indicated by the number in subscript following the drug abbreviation. NT, not tested.

**Table S2:** Change in Q203 plasma concentration after stopping treatment

| Treatment and duration | Mean (±SD) plasma concentration (in $\mu\text{g/ml}$) at the indicated time after the last dose | | |
| --- | --- | --- | --- |
|  | 2-3 days* | 2 weeks | 4 weeks |
| Q<sub>0.5</sub> - 1 week | 0.073 ± 0.024 | 0.008 ± 0.003 | 0.001 ± 0.001 |
| Q<sub>2</sub> - 1 week | 0.104 ± 0.034 | 0.017 ± 0.005 | 0.004 ± 0.0002 |
| Q<sub>10</sub> - 1 week | 0.484 ± 0.100 | 0.095 ± 0.023 | 0.112 ± 0.087 |
| Q<sub>0.5</sub> - 2 weeks | 0.052 ± 0.013 | 0.008 ± 0.002 | 0.002 ± 0.001 |
| Q<sub>2</sub> - 2 weeks | 0.214 ± 0.084 | 0.044 ± 0.029 | 0.005 ± 0.002 |
| Q<sub>10</sub> - 2 weeks | 0.794 ± 0.100 | 0.083 ± 0.023 | 0.021 ± 0.087 |
| R<sub>10</sub>Q<sub>10</sub>- 2 weeks | 0.713 ± 0.052 | 0.226 ± 0.109 | 0.020 ± 0.014 |
| R<sub>20</sub>Q<sub>2</sub>- 2 weeks | 0.204 ± 0.036 | 0.023 ± 0.008 | 0.004 ± 0.001 |
| R<sub>20</sub>Q<sub>10</sub>- 2 weeks | 0.769 ± 0.262 | 0.109 ± 0.060 | 0.011 ± 0.002 |
| *blood draw was 2 days and 3 days after stopping treatment in mice treated for 1 and 2 weeks, respectively |  |  |  |

Drug abbreviations: R, Rifampin; C, Clarithromycin; Q, Q203/Telacebec. Drug dose is indicated by the number in subscript following the drug abbreviation.

**Table S3:** Change in footpad CFU in study 2: *To determine the dose-ranging activity of Q203 alone and in combination with standard and high dose rifampin, including the extended activity after treatment discontinuation*

| Regimen | Mean (±SD) CFU count during treatment | | | | Mean (±SD) CFU count after stopping treatment | | | |
| --- | --- | --- | --- | --- | --- | --- | --- | --- |
|  | D-45 | D0 | Week 1 | Week 2 | Week 1 (+2) | Week 1 (+4) | Week 2 (+2) | Week 2 (+4) |
| Untreated | 2.61 ± 0.21 | 5.4 ± 0.39 | 5.71 ± 0.23 | 5.98 ± 0.23 | - | - | - | - |
| RC | | | 4.79 ± 0.32 | 3.50 ± 0.48 | - | - | 2.61 ± 0.40 | 1.63 ± 0.29 |
| Q<sub>0.5</sub> | | | 5.03 ± 0.43 | 2.24 ± 0.71 | 1.69 ± 0.64 | 0.48 ± 1.48<sup>6</sup> | 0 | 0 |
| Q<sub>2</sub> | | | 3.93 ± 0.58 | 2.00 ± 0.93 | 0 | 0 | 0 | 0 |
| Q<sub>10</sub> | | | 4.25 ± 0.42 | 0.24 ± 0.38<sup>1</sup> | 0.65 ± 0.48<sup>3</sup> | 0 | 0 | 0 |
| R<sub>10</sub> | | | 5.01 ± 0.35 | 4.08 ± 0.41 | 4.81 ± 0.44 | 4.30 ± 1.58 | 2.24 ± 1.40 | 1.24 ± 0.23 |
| R<sub>20</sub> | | | 4.60 ± 0.45 | 4.00 ± 0.49 | 4.87 ± 0.41 | 4.95 ± 1.48 | 1.90 ± 0.55 | 1.78 ± 0.83 |
| R<sub>10</sub>Q<sub>0.5</sub> | | | 4.56 ± 0.73 | 1.87 ± 1.05 | 2.09 ± 1.53<sup>3</sup> | 0.69 ± 0.68<sup>2</sup> | 0.17 ± 0.41<sup>5</sup> | 0.07 ± 0.16<sup>5</sup> |
| R<sub>10</sub>Q<sub>2</sub> | | | 3.98 ± 0.73 | 1.55 ± 1.27 | 0.71 ± 1.00<sup>4</sup> | 0 | 0 | 0 |
| R<sub>10</sub>Q<sub>10</sub> | | | 4.03 ± 0.99 | 1.42 ± 1.57<sup>2</sup> | 1.07 ± 1.29<sup>4</sup> | 0 | 0 | 0 |
| R<sub>20</sub>Q<sub>0.5</sub> | | | 4.45 ± 0.59 | 2.58 ± 0.97 | 1.11 ± 0.83<sup>3</sup> | 0.97 ± 0.53<sup>3</sup> | 0.32 ± 0.28<sup>2</sup> | 0.27 ± 0.30<sup>4</sup> |

<sup>1</sup>1/6 pads negative
<sup>2</sup>2/6 pads negative
<sup>3</sup>1/6 pads negative
<sup>4</sup>3/6 pads negative
<sup>5</sup>5/6 pads negative
<sup>6</sup>CFU count from 3 footpads only, as 3 others were contaminated

Day after infection is D-45. D0 is the start of treatment. Treatment duration is given in weeks and the numbers in parentheses indicate the number of weeks after stopping treatment. Drug abbreviations: R, Rifampin; C, Clarithromycin; Q, Q203/Telacebec. Drugdose is indicated by the number in subscript following the drug abbreviation.

**Table S4:** Experimental scheme for Study 1

| Regimen | Time points for footpad CFU counts |  |  |  |  | Relapse assessment after stopping treatment |  | Total |
| --- | --- | --- | --- | --- | --- | --- | --- | --- |
|  | D-42 | D 0 | Week 1 | Week 2 | Week 4 | Week 2 (+24) | Week 4 (+24) |  |
| <u>Controls</u> |  |  |  |  |  |  |  |  |
| Untreated | 3 (6) | 3 (6) |  | 3 (6) |  |  |  | 9 |
| R<sub>10</sub>CLR<sub>100</sub> |  |  | 3 (6) | 3 (6) | 3 (6) |  | 8 (16) | 17 |
| Q<sub>5</sub> |  |  |  | 3 (6) |  |  |  | 3 |
| Q<sub>10</sub> |  |  | 3 (6) | 3 (6) |  |  |  | 6 |
| <u>Tests</u> |  |  |  |  |  |  |  |  |
| R<sub>10</sub>Q<sub>5</sub> |  |  | 3 (6) | 3 (6) | 3 (6) | 8 (16) | 8 (16) | 25 |
| R<sub>20</sub>Q<sub>5</sub> |  |  | 3 (6) | 3 (6) | 3 (6) | 8 (16) | 8 (16) | 25 |
| R<sub>10</sub>Q<sub>10</sub> |  |  | 3 (6) | 3 (6) | 3 (6) | 8 (16) | 8 (16) | 25 |
| R<sub>20</sub>Q<sub>10</sub> |  |  | 3 (6) | 3 (6) | 3 (6) | 8 (16) | 8 (16) | 25 |
| **Total** | **3 (6)** | **3 (6)** | **18 (36)** | **24 (48)** | **15 (30)** | **32 (64)** | **40 (80)** | **135** |

A total of 135 mice were infected in both hind footpads. At each time point for footpad CFU counts, 3 mice (6 footpads) were harvested. Relapse assessments were done 24 weeks after stopping treatment for 2 or 4 weeks. Drug abbreviations: R, Rifampin; C, Clarithromycin; Q, Q203/Telacebec. Drug dose is indicated by the number in subscript following the drug abbreviation.

**Table S5:** Experimental scheme for Study 2.

| Regimen | Time points for footpad CFU counts during treatment |  |  |  | Time points for CFU counts after stopping treatment |  |  |  | Total mice |
| --- | --- | --- | --- | --- | --- | --- | --- | --- | --- |
|  | D-45 | D0 | Week 1 | Week 2 | Week 1 (+2) | Week 1 (+4) | Week 2 (+2) | Week 2 (+4) |  |
| Untreated | 3 | 3 | 3 (6) | 3 (6) |  |  |  |  | 12 |
| R<sub>10</sub>C<sub>100</sub> |  |  | 3 (6) | 3 (6) |  |  | 3 (6) | 3 (6) | 12 |
| Q<sub>0.5</sub> |  |  | 3 (6) | 3 (6) | 3 (6) | 3 (6) | 3 (6) | 3 (6) | 18 |
| Q<sub>2</sub> |  |  | 3 (6) | 3 (6) | 3 (6) | 3 (6) | 3 (6) | 3 (6) | 18 |
| Q<sub>10</sub> |  |  | 3 (6) | 3 (6) | 3 (6) | 3 (6) | 3 (6) | 3 (6) | 18 |
| R<sub>10</sub> |  |  | 3 (6) | 3 (6) | 3 (6) | 3 (6) | 3 (6) | 3 (6) | 18 |
| R<sub>20</sub> |  |  | 3 (6) | 3 (6) | 3 (6) | 3 (6) | 3 (6) | 3 (6) | 18 |
| R<sub>10</sub>Q<sub>0.5</sub> |  |  | 3 (6) | 3 (6) | 3 (6) | 3 (6) | 3 (6) | 3 (6) | 18 |
| R<sub>10</sub>Q<sub>2</sub> |  |  | 3 (6) | 3 (6) | 3 (6) | 3 (6) | 3 (6) | 3 (6) | 18 |
| R<sub>10</sub>Q<sub>10</sub> |  |  | 3 (6) | 3 (6) | 3 (6) | 3 (6) | 3 (6) | 3 (6) | 18 |
| R<sub>20</sub>Q<sub>0.5</sub> |  |  | 3 (6) | 3 (6) | 3 (6) | 3 (6) | 3 (6) | 3 (6) | 18 |
| R<sub>20</sub>Q<sub>2</sub> |  |  | 3 (6) | 3 (6) | 3 (6) | 3 (6) | 3 (6) | 3 (6) | 18 |
| R<sub>20</sub> Q<sub>10</sub> |  |  | 3 (6) | 3 (6) | 3 (6) | 3 (6) | 3 (6) | 3 (6) | 18 |
| Total | 3 | 3 | 39 (78) | 39 (78) | 33 (66) | 33 (66) | 36 (72) | 36 (72) | 222 |

A total of 222 mice were infected in both hind footpads. At each time point for footpad CFU counts, 3 mice (6 footpads) were harvested. For time points after stopping treatment, (+2) and (+4) indicate time points 2 and 4 weeks, respectively, after completing the indicated duration of treatment. Drug abbreviations: R, Rifampin; C, Clarithromycin; Q, Q203/Telacebec. Drug dose is indicated by the number in subscript following the drug abbreviation.

